# Long read single cell RNA sequencing reveals the isoform diversity of *Plasmodium vivax* transcripts

**DOI:** 10.1101/2022.07.14.500005

**Authors:** Brittany Hazzard, Juliana M. Sá, Angela C. Ellis, Tales V. Pascini, Shuchi Amin, Thomas E. Wellems, David Serre

## Abstract

*Plasmodium* infections often consist of heterogenous populations of parasites at different developmental stages and with distinct transcriptional profiles, which complicates gene expression analyses. The advent of single cell RNA sequencing (scRNA-seq) enabled disentangling this complexity and has provided robust and stage-specific characterization of *Plasmodium* gene expression. However, scRNA-seq information is typically derived from the end of each mRNA molecule (usually the 3’-end) and therefore fails to capture the diversity in transcript isoforms documented in bulk RNA-seq data. Here, we describe the sequencing of scRNA-seq libraries using Pacific Biosciences (PacBio) chemistry to characterize full-length *Plasmodium vivax* transcripts from single cell parasites. Our results show that many *P. vivax* genes are transcribed into multiple isoforms, primarily through variations in untranslated region (UTR) length or splicing, and that the expression of these isoforms is often developmentally regulated. Our findings demonstrate that long read sequencing can be used to characterize mRNA molecules at the single cell level and provides an additional resource to better understand the regulation of gene expression throughout the *Plasmodium* life cycle.

**Author Summary:** Single cell RNA-sequencing is a valuable tool for identifying cell specific differences in heterogenous populations. However, scRNA-seq has limitations in assigning reads to genes of organisims with poorly annotated UTRs, due to the poly-A caputre utilized by some scRNA-seq technologies, this technical limitation also makes identifying transcript specific differences, like alternative splicing, difficult. Despite its importance in human disease the *P. vivax* genome annotation is still relatively sparce, especially in the UTRs, and very little is known about transcript differece in the different life stages of the parasite life cycle. Here, we utilize a modified version of 10X scRNA-seq technology to capture full length transcripts via PacBio sequencing from both sporozoite and blood stages of *P. vivax*. This allowed us to predict full length stage specific transcripts for *P. vivax* as well as identify important variation in the previously poorly annotated UTRs. These findings will aide in futhering our understanding of *P. vivax* transcript regulation across the life cycle stages.

## Introduction

*Plasmodium vivax* is the second most common cause of human malaria worldwide and was responsible for an estimated 4.5 million clinical cases of malaria in 2020(1). Despite these numbers, *P. vivax* research lags behind that of *P. falciparum*, in part due to the difficulty to propagate the parasites *in vitro*(2,3). Genomic techniques, such as genome(4–9) and transcriptome(10–12) sequencing, have improved our knowledge of *P. vivax* biology, but are hindered by the complexity of most blood-stage *P. vivax* infections: multiple genetically-distinct parasites are often simultaneously present in one infection and, due to the lack of, or incomplete, sequestration of *P. vivax* stages, all intraerythrocytic developmental stages concurrently circulate in the blood. Experimental infections of non-human primates using monkey-adapted strains of *P. vivax*(13,14) provide a robust system to study *in vivo* the regulation of all blood stages using monoclonal and well-characterized parasites. However, the simultaneous presence of multiple developmental stages in the blood, each with their own regulatory profiles, remains challenging.

Short-term *ex vivo* cultures(10,11) and statistical inferences of the stage composition(10–12,15) have been used to circumvent this issue but suffer from limited resolution and possible artefacts. Characterization of the gene expression of single cells (scRNA-seq) provides an elegant alternative and has been successfully applied to a wide range of *Plasmodium* species(16–22), including *P. vivax*(23). However, many scRNA-seq assays rely on the capture and sequencing of the 3’ ends of polyadenylated transcripts(24,25) and, consequently, only a short portion of the transcripts is sequenced and the data generated provide little information about transcript isoforms and alternative splicing(11,12,15,26). Additionally, it can be difficult to assign one signal to a specific gene since the scRNA-seq reads often derive from the 3’ untranslated regions (3’-UTRs) which are incompletely annotated in *P. vivax* (11,12,15,23,26).

Here we combine long- and short read sequencing of scRNA-seq libraries to characterize *P. vivax* isoforms throughout the intraerythrocytic life cycle as well as in sporozoites. The standard short read Illumina sequencing of scRNA-seq libraries provides a robust description of the developmental stage of each *P. vivax* parasite captured, while sequencing the same mRNA molecules using PacBio long reads enables characterizing the full-length transcripts present in these cells. The data generated allow to better annotate the 5’- as well as 3’-UTRs of *P. vivax* genes, to comprehensively characterize all transcripts expressed, and to identify stage-specific isoforms. Overall, our results demonstrate that single cell long read sequencing, in conjunction with short read sequencing, provide a robust method for comprehensively characterizing mRNA sequences at the single cell level and identifying isoforms involved in the regulation of specific *Plasmodium* stages.

## Results and Discussion

### Characterization of single cell P. vivax blood-stage and sporozoite transcriptomes

We obtained blood-stage parasites from two *Saimiri boliviensis* monkeys infected with the Chesson strain of *P. vivax* and prepared 10X 3’-end single cell RNA sequencing (scRNA-seq) libraries after enrichment of the infected red blood cells (see *Material and Methods*). After generating 57,550,235-63,399,045 short reads per sample, we successfully mapped 72-74% of all reads to the *P. vivax* P01 genome sequence(27). While full-length mRNAs are captured and converted into cDNA in the 10X droplets, only the 3’-end of each transcript is sequenced due to the cDNA fragmentation occurring during library preparation (**Fig 1**). All 3’-end scRNA-seq reads therefore derive from the last ∼300 bp of the transcripts. However, only 48-52% of the reads mapped within 500 bp of annotated *P. vivax* genes and 47-52% mapped more than 500 bp away from any annotated gene (**Table S1**). These results are consistent with previous analyses(23) and highlight that many genes and/or their 3’-UTRs remain incompletely annotated in the *P. vivax* genome. After removing PCR duplicates and stringent quality control filters, we obtained information about 949 and 1,807 single cell blood-stage transcriptomes, each characterized by more than 5,000 unique reads (**Table 1**). To characterize which blood stages were present in each infection, we analyzed these transcriptomes using principal component analysis and showed that several asexual and sexual developmental stages were present in both samples, although with some differences in their relative proportions (**Fig S1**).

**Figure 1.**
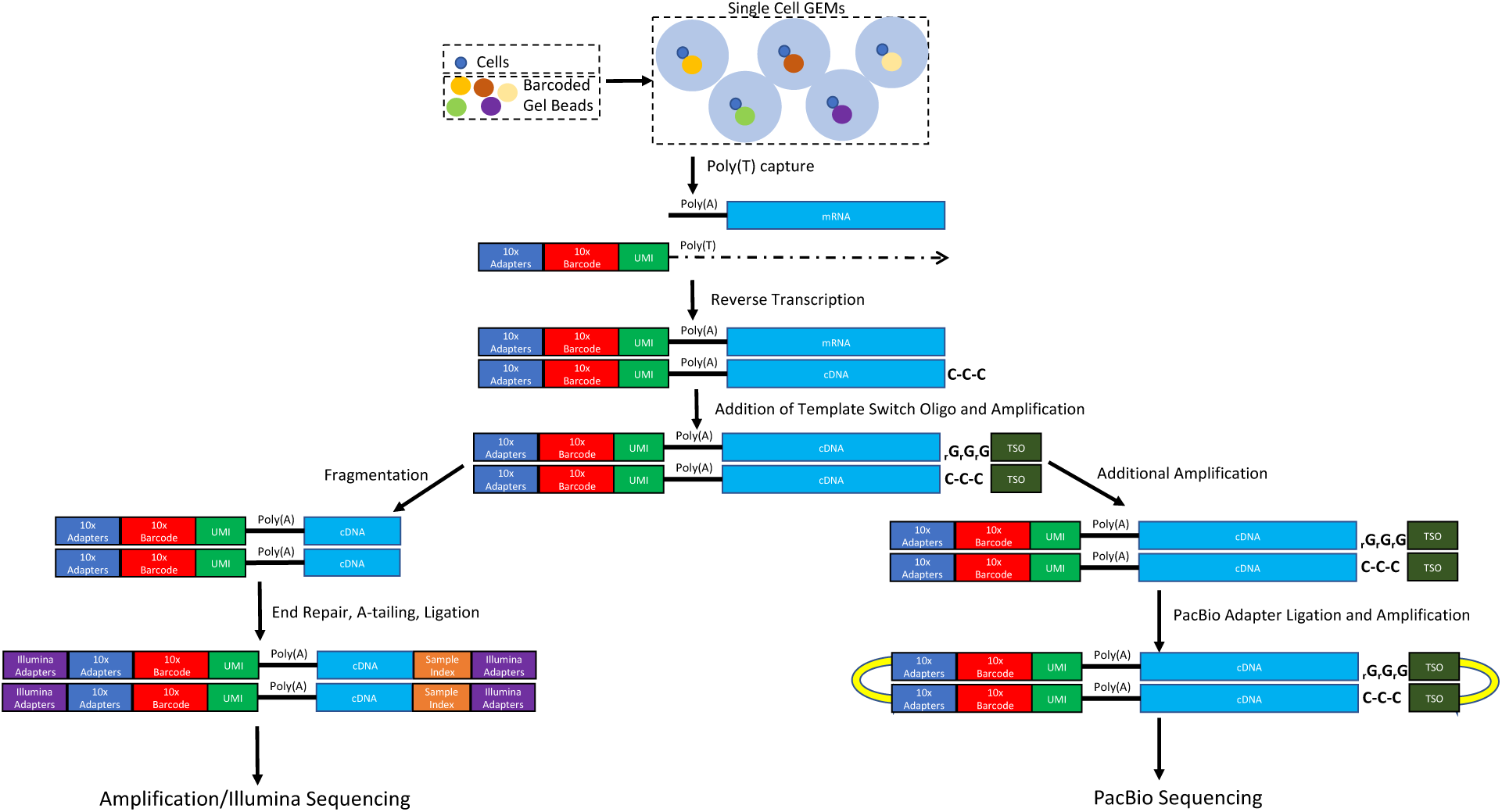
Overview of the experimental design. The figure shows, on the left, the standard 10X scRNA-seq protocol based on Illumina sequencing and, on the right, the protocol used for sequencing full-length transcripts using Pacific Biosciences chemistry. Abbreviations: GEMs, Gel Beads in Emulsion; UMI, Unique Molecule Identifier; TSO, template switch oligo.

**Table 1:** Illumina and PacBio sequencing results and transcript predictions.

| Sample ID | Illumina sequencing data |  |  |  | PacBio sequencing data |  |  |  |
| --- | --- | --- | --- | --- | --- | --- | --- | --- |
|  | Total Reads | %Pv Reads | Unique Reads | No. of Cells* | Total Reads (CCS) | %in Illumina cells | Unique Reads | No. of transcripts |
| Sample 1 (D01) | 57,550,235 | 74% | 31,177,026 | 949 | 3,390,270 | 64% | 1,688,745 | 9,982 |
| Sample 2 (C01) | 63,399,045 | 72% | 35,706,126 | 1807 | 3,570,838 | 50% | 1,523,456 |  |
| Spz 1 | 63,039,836 | 7.08% | 3,654,760 | 2,609 | 2,578,778 | 8% | 106,531 | 784 |
| Spz 2 | 74,606,583 | 3.28% | 2,131,303 | 2,363 | 1,935,640 | 5% | 40,977 |  |

We also prepared two scRNA-seq libraries from salivary gland sporozoites dissected from *Anopheles stephensi* and *Anopheles freeborni*, respectively, fed on *Saimiri* monkeys infected with *P. vivax* parasites. Out of the 63,039,836 and 79,688,059 reads generated from each library, 3-7% mapped to the *P. vivax* genome sequence, similar to the numbers obtained previously for *P. berghei* sporozoites(20). 41-42% of those reads mapped within 500 bp of annotated *P. vivax* genes and 58-59% mapped more than 500 bp away from any gene (**Table S1**). After removing PCR duplicates and stringent quality control filters, we obtained information about 2,609 and 2,363 single cell transcriptomes, each characterized by more than 250 unique reads (**Table 1, Table S1**). We used a lower threshold in this analysis due to the much lower number of *P. vivax* reads recovered as a result of the overwhelming presence of mosquito RNA (but the cutoff is comparable to those previously used for analyzing sporozoite scRNA-seq data(20)). In contrast to the heterogeneity of the blood stage parasites, these salivary gland sporozoites formed a relatively homogeneous population (**Fig S1**).

### Long read sequencing of scRNA-seq libraries provides full-length transcript information

Since only the 3’-end of each transcript is usually sequenced in scRNA-seq experiments, it is sometimes difficult to assign the scRNA-seq reads to a specific gene, or to identify signals derived from different gene isoforms(23). We therefore used cDNA from the same four 10X scRNA-seq libraries before fragmentation to generate full-length isoform sequences using the PacBio chemistry (see **Fig 1** and *Material and Methods* for details).

From each blood sample, we generated 3,390,270 and 3,570,838 circular consensus sequences that each derived from at least 10 passes of sequencing of an individual cDNA molecule. 50-64% of these sequences carried a 10X barcode matching one of the cells characterized by Illumina sequencing, and >99% of those sequences mapped unambiguously to the *P. vivax* genome (**Table 1, Table S1**). After removal of PCR duplicates, we ended up with 1,688,745 and 1,523,456 unique reads (each derived from a unique mRNA molecule) for further analysis (**Table 1**). While the Illumina and PacBio library preparations diverged after cDNA amplification, the same barcoded cDNA molecules were used for both experiments (**Fig 1**) and the numbers of reads obtained with each technology for each individual cell were highly correlated (**Fig S2**).

For the sporozoite samples, we generated 2,578,778 and 1,935,640 circular consensus sequences. 5-8% of these sequences carried a 10X barcode matching one of the cells characterized by Illumina sequencing, and 50-65% of those sequences mapped unambiguously to the *P. vivax* genome. After removal of PCR duplicates and additional QCs, we ended up with 106,531 and 40,977 unique reads respectively for further analysis (**Table 1, Table S1**). Similar to the pattern observed for the blood stage samples, the numbers of reads obtained by Illumina and PacBio sequencing for each sporozoite were also highly correlated (**Fig S2**) although, due to the lower sequencing depth, the overall number of cells with PacBio information was limited to 2,449 and 1,670.

We then summarized these mapped reads into *P. vivax* transcripts and, to avoid including sequences that may represent partially degraded molecules or technical artefacts, we only considered in this analysis predicted transcripts represented by more than 10 PacBio unique reads in both samples of the same type (e.g., in both blood-stage samples). Overall, we identified a total of 9,982 transcripts from blood stage parasites and 784 from sporozoites (**Table 1**), with an average transcript length of 2,432 bp from blood stages and 2,054 bp from sporozoites (ranging from 100 to 20,506 bp and from 197 to 6,809 bp, respectively).

### Most transcripts encode proteins corresponding to the genome annotation

Out of the 9,982 blood stage transcripts, 8,550 transcripts were predicted to encode more than 100 amino acids and 5,368 of those (63%) were predicted to encode a full-length protein (including a start and stop codon) (**Table S1**). The median length of these full-length protein sequences was 291 amino acids, significantly smaller than the length of the protein sequences annotated in the latest version of the P01 *P. vivax* genome (median of 401 amino acids)(27) (**Fig S3**). This discrepancy is likely due to the 10X reverse transcriptase processivity that will hamper synthesis of long cDNA molecules and prevent recovery of full-length transcripts when the mRNA is long (and indeed, most of the full-length transcripts obtained from PacBio sequencing match the annotated reference, see below). Of the 784 sporozoite transcripts, 563 transcripts were predicted to encode more than 100 amino acids and 344 of those (60%) were predicted to encode a full-length protein (including a start and stop codon), with a median length of 267 amino acids (**Fig S3, Table S1**).

We then compared the protein sequences predicted from these full-length transcripts to all *P. vivax* protein sequences annotated in the P01 genome (see **Fig S4**, *Materials and Methods*). Most transcripts were predicted to encode protein-coding sequences highly similar to those annotated in the genome: 4,553 of the blood-stage transcripts (84.8%) and 254 of the sporozoite transcripts (73.8%) were identical to an annotated *P. vivax* protein-coding sequence for >90% of their length (**Fig S5**, see e.g., **Fig 2A**). Additionally, 378 blood-stage (7.1%) and 23 sporozoite transcripts (6.7%) partially matched known protein-coding sequences and were identical over 50%-90% of their sequence. While some of these transcripts could represent instances where the current gene annotation is possibly incorrect (see e.g., **Fig 2B**), in 253 of these cases (63%) another isoform in our dataset matched the entire annotated protein-coding sequence, suggesting that these differences represent incomplete annotations of transcript isoforms rather than incorrect annotations (see also below). One example is the cytochrome b5-like heme/steroid binding protein (PVP01_0716500) that is transcribed as annotated in the genome by blood-stage parasites but, due to an alternative start site, is transcribed into a shorter mRNA by sporozoites, resulting in a shorter predicted protein (**Fig 2C**). Finally, 436 blood-stage (8.1%) and 67 sporozoite transcripts (19.5%) aligned over less than 50% of their length to annotated protein sequences (**Fig S5**) but those transcripts were very short (less than 200 amino acids) and/or had low support from the PacBio reads and likely represented artifacts or fragmented transcripts.

**Figure 2.**
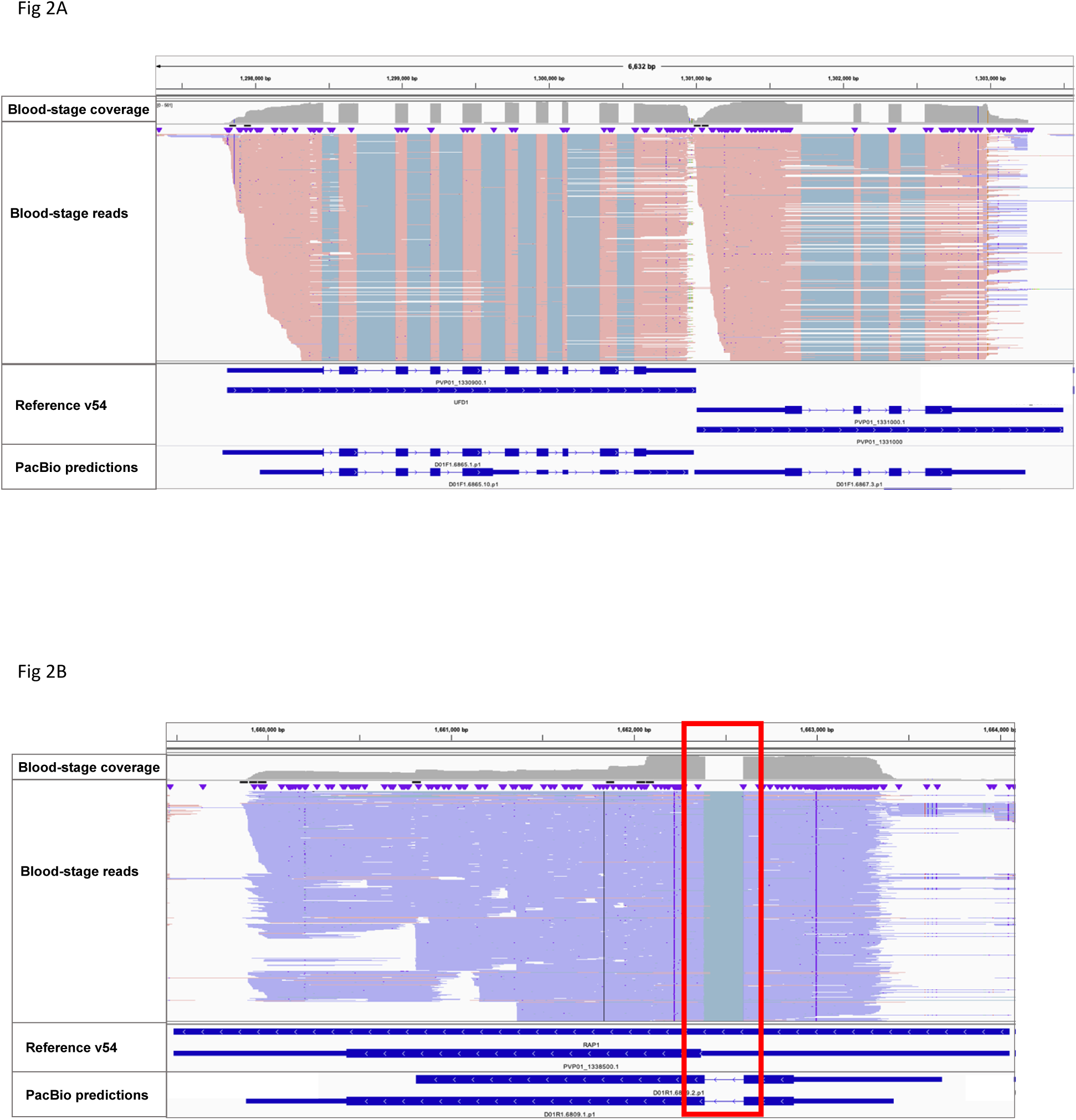
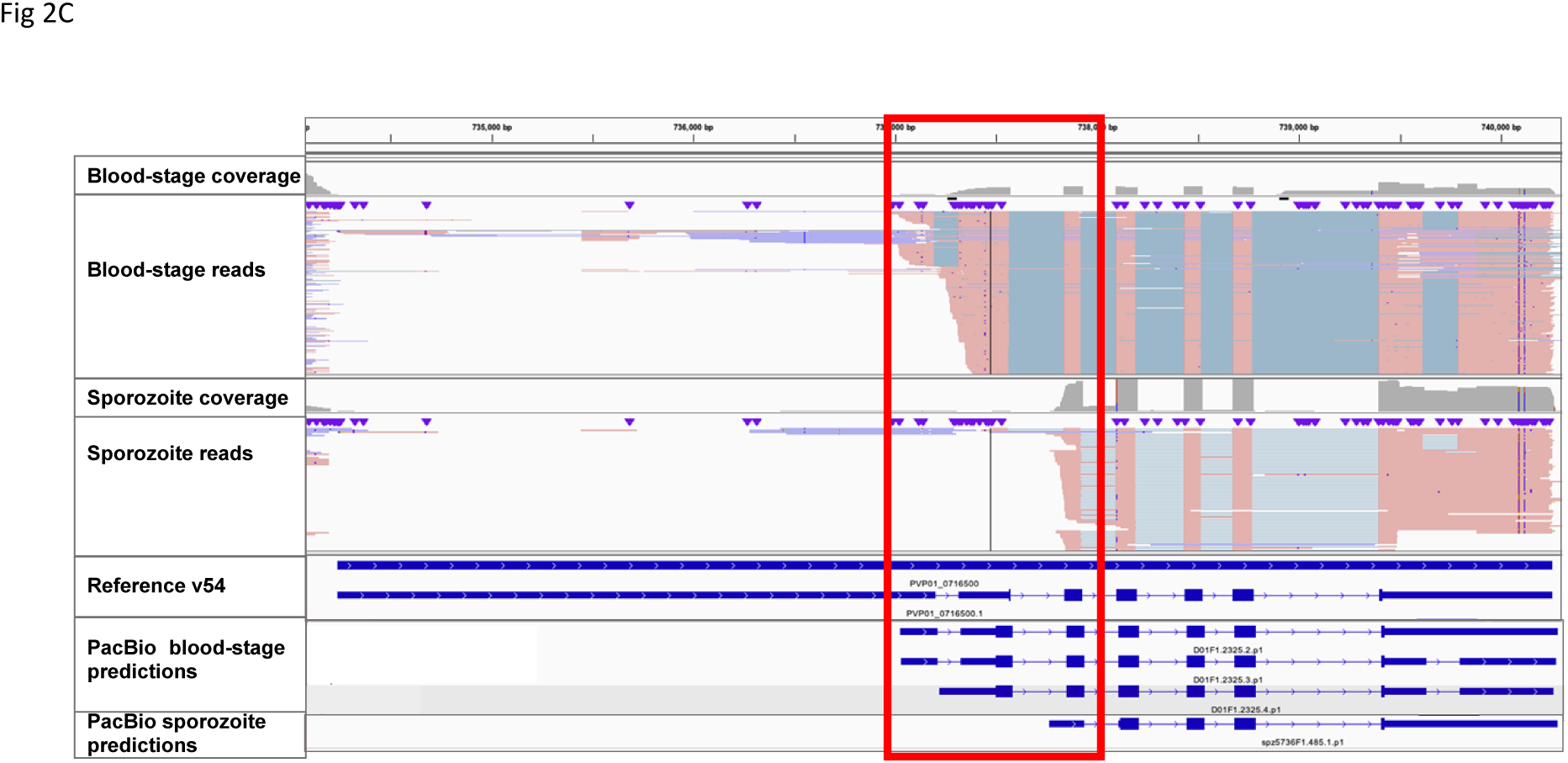
PacBio reads and predicted annotation do not always support the current reference annotation. (**A**) Example of protein-coding transcripts matching the current gene annotations. The figure shows 5.7 kb of chromosome 13 containing two annotated *P. vivax* genes, the ubiquitin fusion degradation protein 1 (PVP01_1330900) and an ATP synthase-associated protein (PVP01_1331000). The blue horizontal bars at the bottom shows the current annotations for these genes in plasmoDB, while the top panel shows the PacBio reads mapping to this locus (each red horizontal line is a unique read mapped to the positive strand with the grey lines indicating spliced introns). Note that while the PacBio reads support a shorter 3’-UTR than annotated for PVP01_133100 the predicted protein coding sequence is identical to the one annotated. **(B)** Example of protein coding transcript differing from the current gene annotation. The figure shows 5 kb of chromosome 13 surrounding the rhoptry-assicated protein 1 (RAP1, PVP01_1338500). The PacBio reads (mapped to the negative strand and displayed in blue) support the presence of an unannotated intron (red box), leading to additional predicted coding sequences upstream of this intron and a different protein than annotated in the genome (thick blue bars at the bottom). **(C)** Example of two isoforms with different predicted protein coding sequences. The top panel shows that the blood-stage parasites express a transcript for cytochrome b5-like heme/steroid binding protein (PVP01_0716500) identical to the annotated protein-coding sequence (although with a shorter 5’-UTR). The middle panel shows that *P. vivax* sporozoites express this gene from a different start site (red box) resulting in a shorter transcript and a different predicted protein.

### Most P. vivax transcripts have extended UTRs that often contain introns

Even for the transcripts highly similar to the annotated protein-coding genes, the PacBio sequences add novel information by providing a detailed description of the 5’- and 3’-UTRs which, despite recent efforts(26), remain incompletely annotated in *P. vivax*. Consistent with previous reports(11,15), our data showed extensive UTRs in many genes (**Fig 3**). We observed that 5’- and 3’-UTRs were roughly similar in length in transcripts expressed by blood-stage parasites, while 5’-UTRs were, on average, slightly longer than 3’-UTRs in sporozoite transcripts (745 vs 731 bp in blood stages [p-value = 0.3149] and 762 vs 650 bp in sporozoites [p-value = 0.0076]). While they have not been extensively studied in *Plasmodium*, in many eukaryotes UTRs contain important regulatory elements which often affect mRNA stability(28–32). While the length of the 5’-UTR does not appear to be associated with the level of the transcript expression determined using the short reads (p=0.315), the 3’-UTR length was negatively correlated with expression level (p=2.8×10^−6^), although with a very low coefficient of correlation (Spearman’s Rho = -0.0798) (**Fig S6**). In an attempt to identify regulatory elements in the UTRs, we compared the abundance of all possible 5-mers in the 5’-UTRs, 3’-UTRs and promoter regions but failed to detect significant enrichment (**Fig S7**). More in-depth analyses will be required to understand the function of these UTRs but the data presented here will provide a solid foundation to implement these studies.

**Figure 3.**
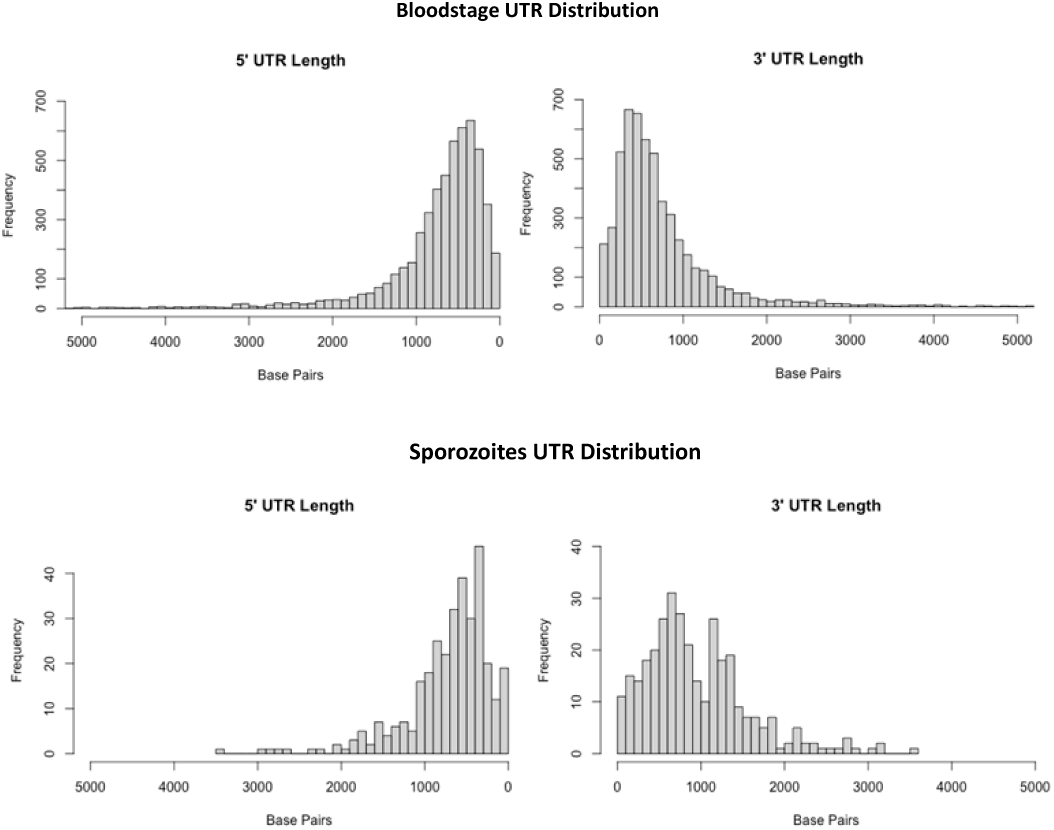
UTR length distributions. Distribution of UTR lengths for transcripts expressed by blood-stage parasites (top) and sporozoites (bottom).

Interestingly, out of the 5,368 full-length blood stage protein-coding transcripts, 1,072 (20%) had at least one intron in the UTRs: 419 transcripts had at least one intron in the 5’-UTR, 568 in the 3’-UTR and 89 had introns in both UTRs (see e.g., **Fig 4**). Of these, 322 transcripts had more than one intron in their UTRs. Of the 344 full-length sporozoite transcripts, 26 had introns in the UTRs, 12 had at least one intron in the 5’-UTR, 11 in the 3’-UTR and 3 had introns in both UTRs. Five of these transcripts had more than one intron in the 5’- or 3’-UTR. Remarkably, in blood-stage parasites the transcripts of genes with an intron in the UTR were, on average, 3-fold higher than those of genes with no UTR intron (Wilcoxon rank test, p-value < 2.2×10^−16^), suggesting that presence of introns in the UTR may be associated with increased mRNA stability in *P. vivax*. Furthermore, genes with introns in their coding sequences were also three times more likely to have introns in the UTR (χ^2^ = 206.78, p-value < 2.2e-16), which may suggest that UTR splicing is mechanistically associated with splicing of coding sequences, possibly due to a more effective recruitment of the splicing machinery at those transcripts. To evaluate whether the presence of UTR introns was more frequent, at a specific developmental stage, we assigned each gene to the blood stage where it was most abundantly expressed. The proportion of genes with introns in their UTR was significantly different among stages (χ^2^= 26.141, p-value = 8.9 × 10^−6^), with 11% and 18% of the genes most expressed in early and late trophozoites having an UTR intron, respectively, while only 3% and 8% of the genes expressed in female gametocytes and schizonts, respectively, did (note that genes with multiple isoforms were excluded from this analysis as it is difficult to robustly quantify the relative expression of isoforms). Finally, we tested whether genes containing introns in their UTRs disproportionally belonged to a specific pathway but failed to detect any gene ontology enrichment (FDR <= 0.1, **Table S3**).

**Figure 4.**
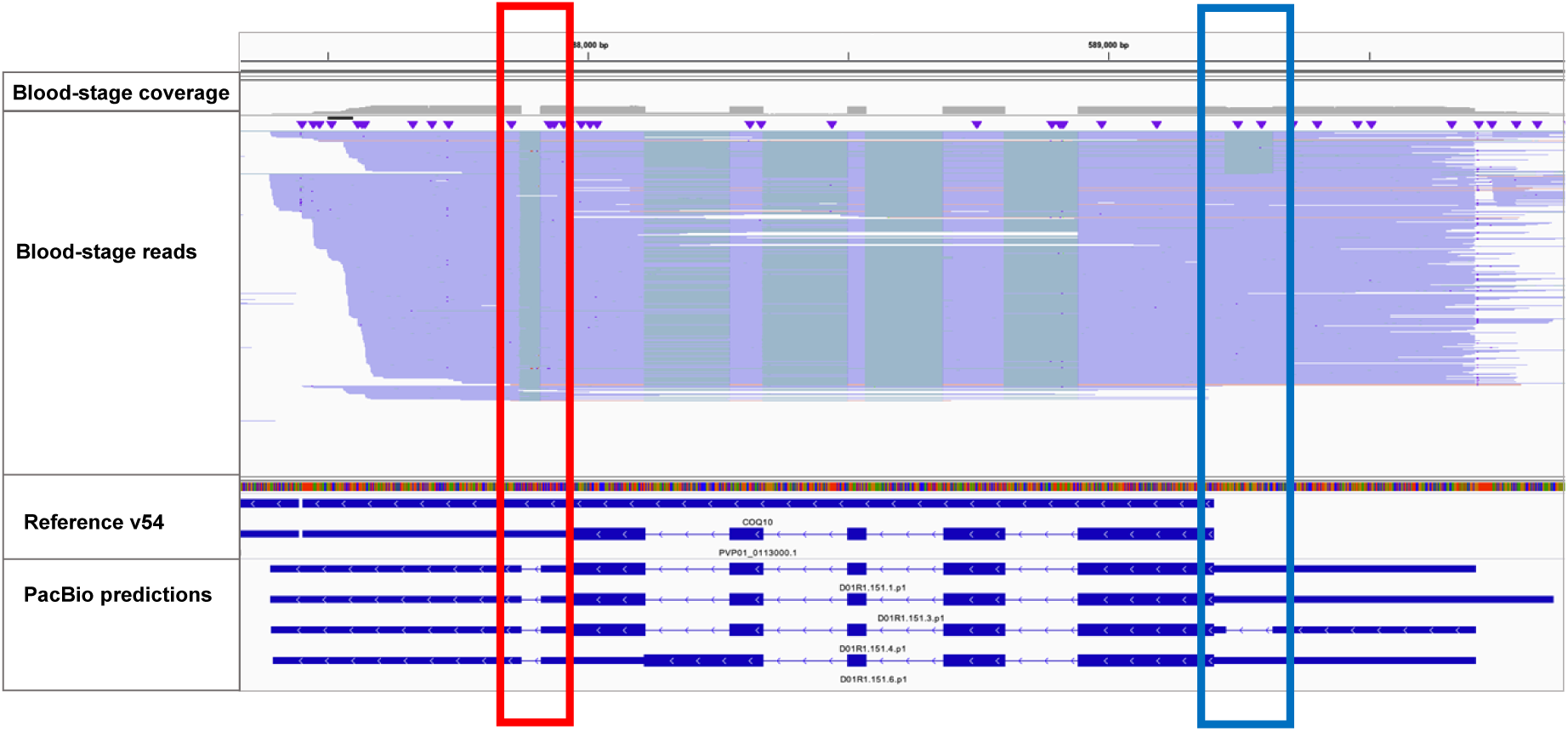
Example of a transcript with unannotated UTR introns. The figure shows PacBio reads (in blue) corresponding to the annotated mRNA for coenzyme Q-binding protein COQ10 homolog (PVP01_0113000) but with an unannotated intron in the 3’-UTR (red box) as well as, for a subset of the mRNAs, a second intron in the 5’-UTR (blue box).

### Transcript isoforms are common in P. vivax and can be expressed in a stage-specific manner

The 5,368 full-length protein coding transcripts derived from blood-stage parasites were transcribed from 2,869 genes: 1,687 genes (59%) were transcribed into a single isoform, while 1,182 (41%) showed evidence of multiple isoforms (and out of those, 719 genes were expressed in two isoforms, 247 in three and 216 were transcribed in four or more isoforms). Most isoforms (n= 819, 70%) encoded the same protein sequence and differed only in their UTR length: 1,024 (87%) genes with isoforms differed in their 5’-UTR length and 1,008 (85%) differed in the 3’-UTR length, with 450 genes with isoforms differing in their UTR introns (**Table 2**, see also **Fig 4** for an example). 363 genes showed evidence of isoforms that were predicted to encode for different proteins, due to an alternate protein coding start (299), alternative end (311), and/or exon skipping (67) (**Table 2**).

**Table 2.**
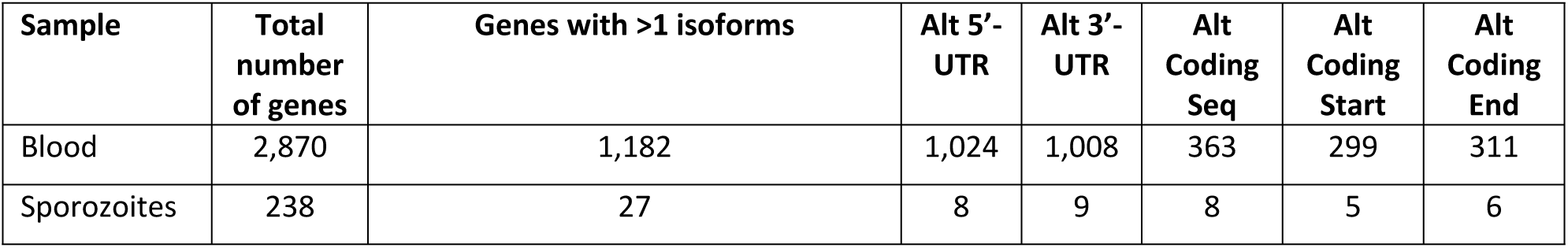
Summary of isoform types from PacBio predictions.

| Sample | Total number of genes | Genes with >1 isoforms | Alt 5'-UTR | Alt 3'-UTR | Alt Coding Seq | Alt Coding Start | Alt Coding End |
| --- | --- | --- | --- | --- | --- | --- | --- |
| Blood | 2,870 | 1,182 | 1,024 | 1,008 | 363 | 299 | 311 |
| Sporozoites | 238 | 27 | 8 | 9 | 8 | 5 | 6 |

In sporozoites, the 344 full-length protein coding transcripts were transcribed from 238 genes: 211 genes (89%) were transcribed into a single isoform, while 27 (11%) showed evidence of multiple isoforms (20 genes were expressed in two isoforms and three in three isoforms and four in four or more). All but eight isoforms were predicted to encode the same protein. Five of these genes had an alternate coding start and six had an alternate coding stop resulting in the change in coding sequence, and all had an exon skip or truncation (**Table 2**).

Since the full-length transcript data derived from molecules characterized by scRNA-seq, these data provide a unique opportunity to preliminarily examine whether different isoforms were expressed at different stages of the parasite development. Using the 10X cell barcodes, we determined the developmental age of each cell using pseudotime determined from the Illumina scRNA-seq data. We only considered in this analysis the 636 genes that had two (or more) isoforms expressed in at least 50 individual cells each. 123 (19%) of these genes showed evidence of expressing isoforms according to the parasite development (**Table S4**). These stage-specific isoforms included 62 and 35 genes with differences in 5’- or 3’-UTR length (see e.g., **Fig 5A** or **Fig 5B**) and/or changes in coding sequences (n=40) (**Fig 5C**). These findings suggest that *P. vivax* utilizes alternative start or termination of transcription as a means of transcriptional or translational regulation between stages.

**Figure 5.**
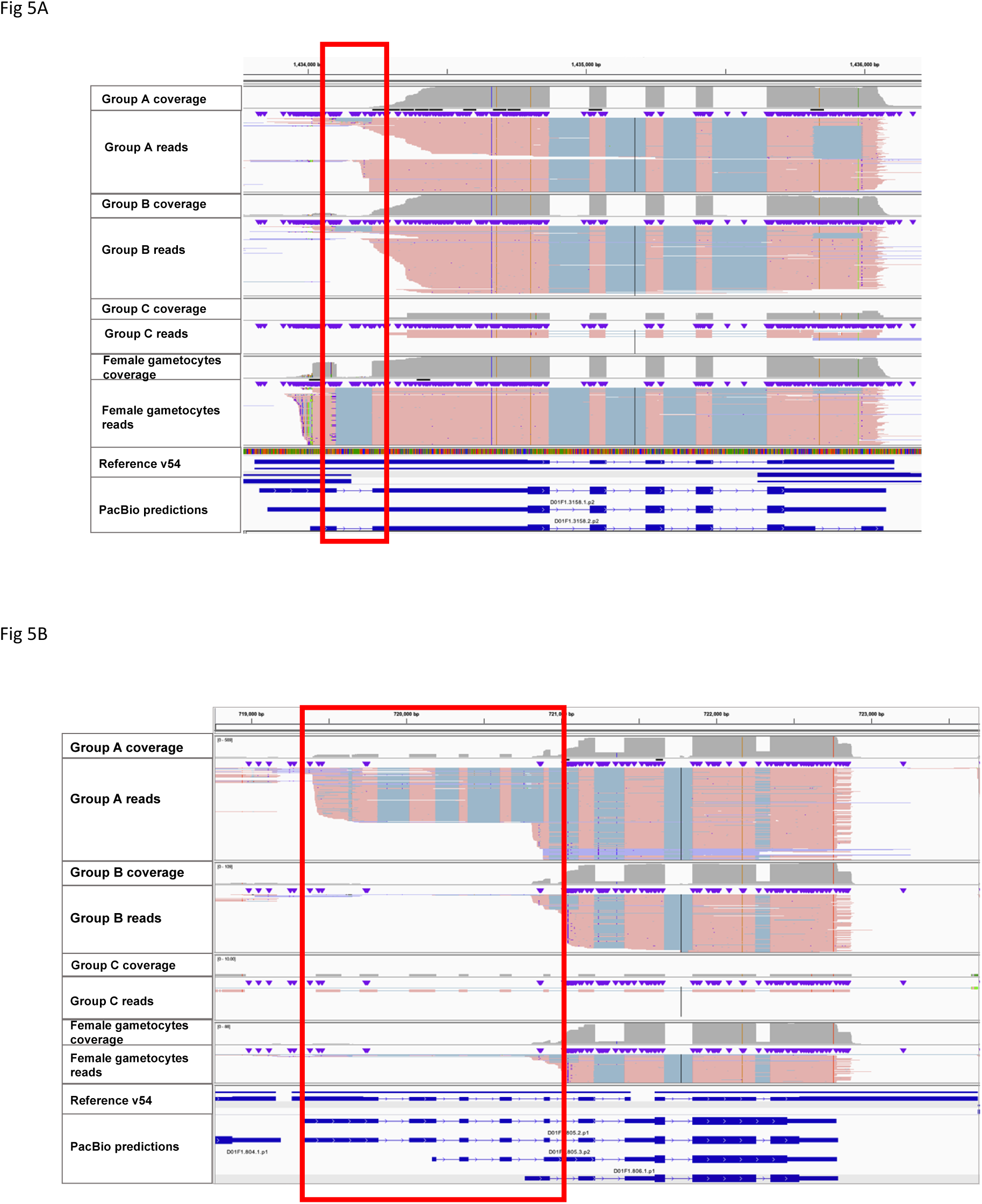
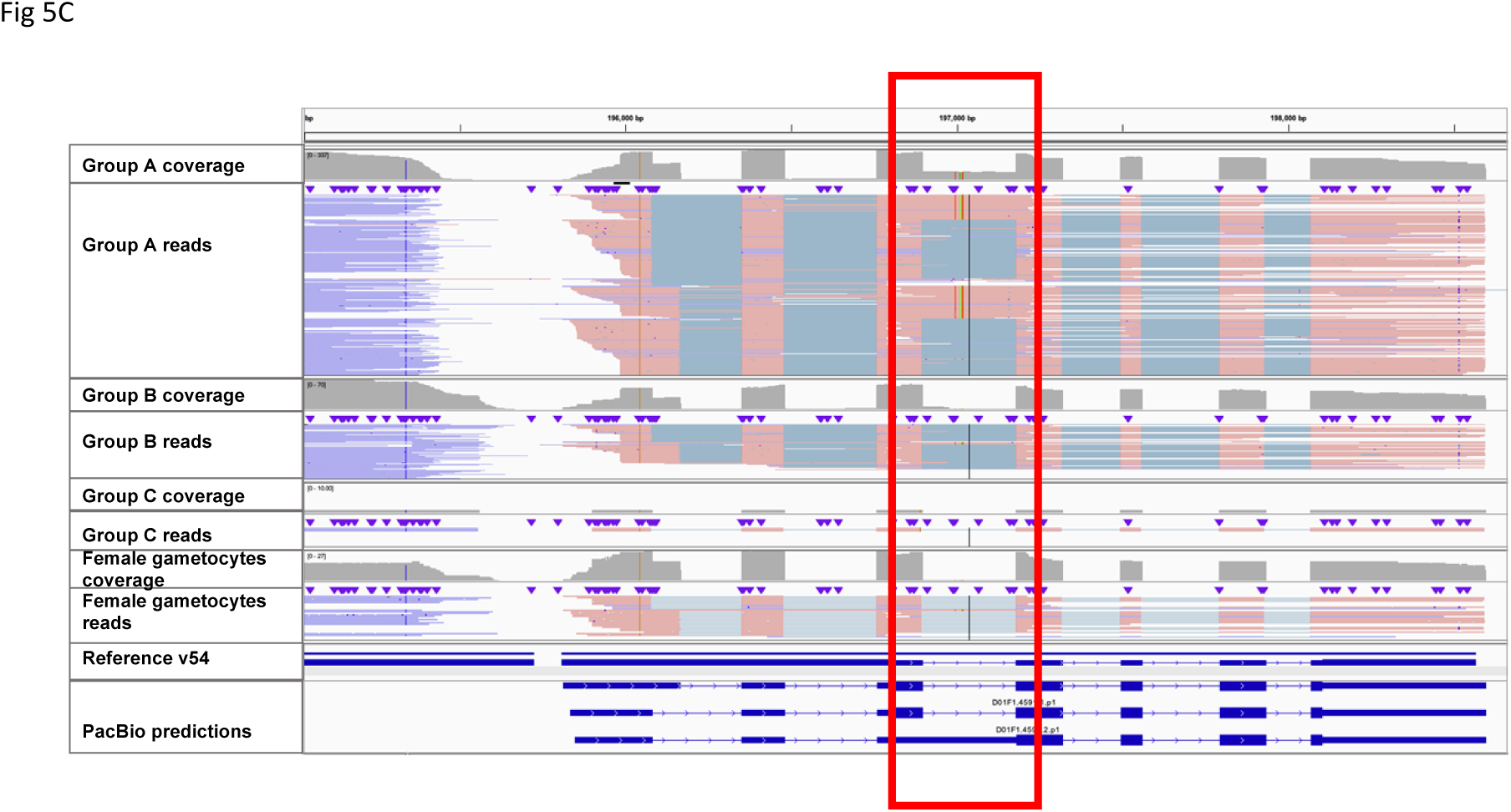
Examples of isoforms expressed in a stage-specific manner. Each panel shows the PacBio reads mapped to a selected locus and split in four groups according to the stage of the parasites they derived from: early trophozoites (group A), late trophozoites (group B), schizonts (group C) and female gametocytes (from top to bottom). **(A)** Female gametocytes express glutaredoxin 1 (PVP01_0833900) from a more upstream TSS than asexual parasites, and the resulting transcripts have a intron in the 5’-UTR (red box). (**B**) Early trophozoites express the Ham 1-like protein (PVP01_0316500) from a more upstream TSS than the other stages and the resulting transcripts have a longer 5’-UTR containing five introns (red box). **(C)** In contrast to the examples shown in **(A)** and **(B)**, the isoforms expressed from suppressor of kinetochore protein 1 (PVP01_1105000) result into different predicted protein coding sequences: some transcripts expressed exclusively in early trophozoites retain the third intron (red box) leading to a different open reading frame (blue bars at the bottom).

### Conclusion

Long read PacBio sequencing coupled with short read Illumina sequencing of single cell RNA-seq libraries allows detection and characterization of full-length transcripts as well as analysis of mRNA isoforms according to the parasites’ developmental stages. Our study highlights the presence of extensive, and incompletely unannotated, UTRs and of UTR introns, that are sometimes expressed in a stage-specific manner. These results suggest that non-coding regions may influence the regulation of gene expression in *P. vivax* and provide a solid framework to further investigate this mechanism.

## Materials and Methods

### Ethics statement

All animal procedures were conducted in accordance with the National Institutes of Health (NIH) guidelines and regulations(33), under approved protocols by the National Institute of Allergy and Infectious Diseases (NIAID) Animal Care and Use Committee (ACUC) (Animal study NIAID LMVR15). Animals were purchased from NIH-approved sources and transported and housed according to Guide for the Care and Use of Laboratory Animals(33).

### Animal studies and sample collection

We infected two splenectomized *Saimiri boliviensis* monkeys with the Chesson strain of *P. vivax* using parasitized erythrocytes from cryopreserved stocks. Once they developed a parasitemia >0.1%, we collected 1 mL of blood sample from the femoral vein of each monkey after anesthesia with 10 mg/kg of ketamine and processed the blood samples on MACS LS columns as previously described(23).

Two blood samples from two additional *Saimiri boliviensis* monkeys coinfected with a NIH-1993 clone(23) nd the Chesson *P. vivax* strain were used for membrane feeding of *Anopheles stephensi* and *Anopheles freeborni*. Salivary glands sporozoites were then collected from each feeding at 21 days post-feed: 50 female mosquitoes were anesthetized on ice and their salivary glands dissected in PBS under a stereomicroscope. The salivary glands were transferred to a low-retention tube (Protein LoBind Tube; Eppendorf) containing PBS, homogenized with a disposable pestle, span down, washed, resuspended, and quantified.

### 10X single cell RNA-sequencing library preparation and sequencing

An estimated 3,000 infected red blood cells or sporozoites from each sample were loaded onto 10X Chromium controller to prepare scRNA-seq libraries according to the manufacturer’s instructions. We then generated, from each library, 57-75 million paired-end reads using an Illumina NovaSeq 4000.

In addition, an aliquot of the cDNA prior to fragmentation was amplified by eight additional cycles of PCR before preparation of a PacBio library using the SMRTBell Express kit 2.0. We then generated 196-328 million reads from each library using a PacBio Sequel II.

### Short read analysis and single cell characterization

Following Illumina sequencing, the short reads were processed as described previously(23). Briefly, we mapped all reads to the P01 *P. vivax* genome(27) using Hisat2(34) with the default parameters except for a maximum intron length of 5,000 bp. We then removed PCR duplicates by identifying reads with identical barcode, unique transcript identifier (UMI) and mapping coordinates. We assigned each unique read to i) a cell based on its barcode and ii) a 500 bp window based on its genomic position. Only cells defined by more than 5,000 unique reads in blood stages and 250 unique reads in sporozoites were further analyzed. Count matrixes and principal component analysis was performed via in-house scripts as well as scran(35).

### Long read analysis

The entire analytical pipeline for processing and analyzing the long-read data is described in **Fig S4**. Briefly, the raw PacBio reads were first collapsed into circular consensus sequencing (CCS) using smrtanalysis(36) and only CCS supported by more than 10 passes were considered for further analysis. We then compared the 10X barcodes of each CCS reads with those obtained after Illumina sequencing and kept all reads matching the barcodes of one of the cells characterized by more than 5,000 unique Illumina reads. To account for the higher error rate of PacBio sequencing we allowed up to one nucleotide mismatch in the barcode sequence. After trimming the 10X adapters, 10X barcodes and the polyA tails, we mapped all CCS reads to the *P. vivax* P01 genome using minimap2(37) using the cdna parameters and a *k* of 14,. PCR duplicates were removed as described above. Only reads mapped to the P01 genome over 50% of their length were kept for further analysis.

### Transcript and protein identification

Transcript prediction was performed separately for each sample utilizing Stringtie2(38,39). Mapping files were divided by forward and reverse reads and single and multiple exon reads. Each of the four files was run separately using Stringtie2 long read default parameters. Only transcripts supported by more than 10 reads in each sample were considered for further processing (separately for the blood-stage and sporozoite samples). Transcript predictions were then compared and collapsed across samples into a single gtf for the blood-stage and sporozoite samples using gffcompare and custom scripts. We then used Transdecoder(40) to identify putative protein coding sequences from the predicted transcript sequences. Finally, we compared the predicted protein sequences to those annotated in the most current *P. vivax* P01 genome (v54) using BlastP and custom scripts.

### Stage-specific transcript analysis

To identify isoforms expressed in a stage-specific manner, we first assigned each PacBio read to a specific isoform using BLAT to against all predicted stringtie transcripts (including non-coding and incomplete transcripts) and considering the match with the greatest overall identity (and only considering reads aligned with >90% identity as aligned). We then identify the pseudotime of the cell expressing this transcript by matching the GEMS barcode to the Illumina data and tested differences in the pseudotime of the cells expressing each isoform of the same gene using a Two-sample Kolmogorov–Smirnov test.

## Data Availability

All sequence data generated in this study are deposited at the Sequence Read Archive under the BioProject XXXX. Custom scripts are available through github at https://github.com/bhazzard11/Single-Cell-PacBio.

## Acknowledgments

We thank the technicians, care-takers, and veterinaries of the Division of Veterinary Resources and of the insectary of the Laboratory of Malaria and Vector Research, National Institute of Allergy and Infectious Diseases, for animal care, technical assistance, and mosquito rearing; and S. Ott, H. Bowen, L. Sadzewicz and L. Tallon in the Genomic Resource Center at the University of Maryland School of Medicine for their support with Illumina and PacBio sequencing.

## Supplemental Figure legends

**Figure S1. Principal components analysis showing the relationship among the individual parasite cells characterized by scRNA-seq (using the Illumina data).** Top row: each dot is a single cell blood stage transcriptome and is displayed based on it gene expression profile and colored according to the expression of stage markers: red – early trophozoites, green – late trophozoites, purple – schizonts, turquoise – female gametocytes. Bottom row: single cell sporozoite data.

**Figure S2. Illumina and PacBio reads correlation.** Correlation between the number of Illumina reads (x-axis) and PacBio reads (y-axis) obtained from each cell (individual black dots).

**Figure S3. Predicted protein length distributions.** Distribution of the length (in amino acids) of the protein-coding sequences predicted from the PacBio transcripts (in red) and of the protein-coding sequences currently annotated in the P01 *P. vivax* genome (in blue). (Note that the x-axis is cut and extends to 12,000 in both panels).

**Figure S4. Summary of the bioinformatic pipelines used for processing the PacBio reads and predicting transcripts.**

**Figure S5. Predicted protein comparision to reference.** Distribution of the percentage alignment (x-axis) of the predicted protein coding sequences with the most similar protein sequence annotated in the P01 genome. Left: blood-stage transcript, right: sporozoite transcripts. (Note that the y-axis is cut and the right-most bars go to 400 and 250 for the left and right panels, respectively).

**Figure S6. Predicted UTR length analysis.** Correlation between the length of a transcript’s UTR (x-axis) and its level of expression determined by Illumina data (y-axis). Note that only genes expressing a single isoform are included in this analysis.

**Figure S7. UTR kmer anaylsis.** Comparison of the abundance of all 5-mers in gene promoters, 5’-UTRs and 3’-UTRs.

**Table S1. Sequencing and transcript predictions results.** Table listing numbers from each step of sequencing analysis and transcript predictions.

**Table S2. Descript list of predicted transcripts.** Table listing all predicted transcripts, there nucelotide sequence, amino acid sequence (if applicable), PVP01 gene name and discription (if applicable), UTR lengths, and number of introns.

**Table S3. Results of Gene Ontology analysis.** Top hits from GO analysis.

**Table S4. Descriptive list of predicted genes with isoforms.** List of PacBio transctipts determined to have isoforms, there PVP01 reference name and desctriptsion, number of isoforms and type of isoform (coding or UTR).

